# Gene expression and chromatin accessibility comparison in iPSC-derived microglia in African, European, and Amerindian genomes in Alzheimer’s patients and controls

**DOI:** 10.1101/2024.08.27.609943

**Authors:** Sofia Moura, Luciana Bertholim Nasciben, Aura M. Ramirez, Lauren Coombs, Joe Rivero, Derek J. Van Booven, Brooke A. DeRosa, Kara L. Hamilton-Nelson, Patrice L. Whitehead, Larry D. Adams, Takiyah D. Starks, Pedro R. Mena, Maryenela Illanes-Manrique, Sergio Tejada, Goldie S. Byrd, Mario R. Cornejo-Olivas, Briseida E. Feliciano-Astacio, Karen Nuytemans, Liyong Wang, Margaret A. Pericak-Vance, Derek M. Dykxhoorn, Farid Rajabli, Anthony J. Griswold, Juan I. Young, Jeffery M. Vance

**Affiliations:** John P. Hussman Institute for Human Genomics, University of Miami Miller School of Medicine, Miami, Florida, USA; Maya Angelou Center for Health Equity, Wake Forest University, Winston-Salem, North Carolina, USA; Neurogenetics Working Group, Universidad Científica del Sur, Lima, Peru; Neurogenetics Research Center, Instituto Nacional de Ciencias Neurológicas, Lima, Peru; Universidad Central del Caribe, Bayamon, Puerto Rico, USA; Dr. John T. Macdonald Foundation Department of Human Genetics, University of Miami Miller School of Medicine, Miami, FL, USA

**Author notes:** Correspondence to Jeffery M. Vance.

## Abstract

Alzheimer’s disease (AD) risk differs between population groups, with African Americans and Hispanics being the most affected groups compared to non-Hispanic Whites. Genetic factors contribute significant risk to AD, but the genetic regulatory architectures (GRA) have primarily been studied in Europeans. Many AD genes are expressed in microglia; thus, we explored the impact of genetic ancestry (Amerindian (AI), African (AF), and European (EU)) on the GRA in iPSC-derived microglia from 13 individuals (∼4 each with high global ancestry, AD and controls) through ATAC-seq and RNA-seq analyses. We identified several differentially accessible and expressed genes (2 and 10 AD-related, respectively) between ancestry groups. We also found a high correlation between the transcriptomes of iPSC-derived and brain microglia, supporting their use in human studies. This study provides valuable insights into genetically diverse microglia beyond the analysis of AD.

## Introduction

Alzheimer’s Disease (AD) affects millions of people worldwide with currently ∼11% of the US population (65 and older) affected. It is predicted that over 150 million individuals will be affected by AD worldwide by 2050. Pathologically, AD is characterized by β-amyloid (Aβ) deposition as neuritic plaques and intracellular accumulation of hyperphosphorylated tau as neurofibrillary tangles, all of which lead to neurodegeneration and progressive cognitive impairment ^1^.

African American (AA) and Hispanic (HI) individuals have the highest risk of developing AD, followed by non-Hispanic White (NHW) individuals, likely due to a combination of environmental and genetic factors. Specifically, in the US, AD affects 19% of AA, 14% of Hispanics, and 10% of NHW ^2^. Further, over the next 25 years, the greatest growth in AD will be in Africa and South America. Genetic diversity and admixture play important roles in disease risk. African American genomes are typically admixed between African and European ancestries while HI encompass a three-way admixture of European, Amerindian, and African ancestries ^3^. Consistent with this, there are ancestry-related differences in the genetic architecture of AD ^4^. Although there are gene variants consistently associated with AD risk across different populations, recent genome-wide association studies (GWAS) have identified several ancestry-specific risk variants, including variants in *ABCA7* ^5–9^, *MPDZ* ^10^, and *IGF1R* ^5,10^. Thus, it is crucial to investigate ancestry-specific disease mechanisms to understand the differential disease susceptibility in different populations and to facilitate the move toward personalized medicine across ancestries.

Most AD-associated and GWAS ^11–13^ risk loci lie in non-coding, regulatory regions. However, the regulatory architecture of the genome has not been extensively analyzed in diverse populations, with most of the existing data derived from individuals of European ancestry. The different population risk profiles for AD of *APOEe4* carriers of different ancestry present a clear example of how differences in gene regulation can affect AD susceptibility. Rajabli *et al.* demonstrated that the lower risk for AD in carriers of *APOEe4* with African ancestry relative to European ancestry was due to differences in the local genomic ancestry surrounding the *APOEe4* allele ^14,15^. Subsequently, it was found that European local genomic ancestry carriers of *APOEe4* had higher *APOEe4* expression and more open chromatin accessibility than that of African local ancestry carriers ^16,17^, supporting the recent report that lower expression of *APOEe4* is tied to lower risk ^18^ and highlighting ancestral differences in gene regulatory networks.

Although much of AD pathogenesis research has focused primarily on neurons, studies suggest a critical role for microglia in the AD disease process. Autopsy studies found an elevated proportion of activated microglia significantly correlated with pathological AD ^19^, specifically the total Aβ load and number of neuritic plaques. Furthermore, a large number of reported AD GWAS genes are expressed in microglia ^20,21^, further supporting their role in AD pathology. Microglia are the resident immune cells of the central nervous system (CNS) and play key roles in brain development, synaptic pruning, homeostasis, and neuronal network maintenance, among other immune response processes ^22^. Specifically, in the context of AD, microglia are particularly important for Aβ plaque clearance, neuroprotection, inflammatory responses, and synaptic homeostasis ^23^.

Here we report an examination of iPSC-derived microglia from African, European, and Amerindian ancestries, expanding on our previous studies of single nuclei RNA-seq and single nuclei ATAC-seq on postmortem microglia from the frontal cortex on African and European genomes ^16,17^. Additionally, as iPSC-derived cells have become important models for human neurodegenerative research, we performed a comparison between our iPSC-derived microglia and autopsy samples to determine similarities and differences. While this study is focused on AD-GWAS genes, this data will be useful for all neurological genetic studies of African, European, and Amerindian populations, as well as admixed populations of African American and Hispanic individuals.

## Results

### Differentiation and validation of iPSC-derived microglia

We differentiated thirteen iPSC-derived microglia (iMGL) lines from individuals of diverse ancestral backgrounds, AD cases and controls, males and females, all derived from individuals over 65 years of age (**Table 1**). Specifically, we differentiated 4 Amerindian (AI), 5 European (EU), and 4 African (AF) iMGL lines. Genotyping and whole genome sequencing were performed to 1) identify the global ancestry and 2) confirm the absence of known mutations in AD-related Mendelian genes (*APP, ABCA7, MAPT, PSEN1, PSEN2, SORL1,* and *TREM2*; **Supplementary Table 1**) that could affect the GRA.

**Table 1:** iPSC-derived Microglia cell line information. AI: Amerindian. EU: European. AF: African. AD: Alzheimer’s disease. MCI: Mild Cognitive Impairment.

| Sample | Global Ancestry |  | Age | Sex | APOE | Clinical Diagnosis |
| --- | --- | --- | --- | --- | --- | --- |
| 1 | AI | 96.3% | 86 | Male | 3/3 | Control |
| 2 | AI | 95.5% | 86 | Male | 3/3 | Control |
| 3 | AI | 100% | 71 | Female | 4/4 | AD |
| 4 | AI | 92.0% | 86 | Female | 3/3 | Control |
| 5 | EU | 100.0% | 88 | Male | 4/4 | AD |
| 6 | EU | 88.6% | 76 | Female | 4/4 | AD |
| 7 | EU | 99.7% | 65 | Female | 3/3 | Control |
| 8 | EU | 93.8% | 67 | Female | 3/3 | Control |
| 9 | EU | 99.5% | 72 | Female | 4/4 | AD |
| 10 | AF | 94.4% | 70 | Female | 4/4 | AD |
| 11 | AF | 96.4% | 75 | Female | 3/3 | Control |
| 12 | AF | 91.5% | 84 | Female | 3/3 | AD |
| 13 | AF | 93.5% | 90 | Female | 3/3 | MCI |

All thirteen iMGL cell lines were further validated with microglia cell-specific lineage markers using immunocytochemistry (ICC) (*PU.1* (*SPI1*)*, TMEM119, TREM2, and P2RY12*; **Supplementary Figure 1**). All microglia cell lines expressed these cell-type specific markers, and their transcriptomic profiles correlated well (r=0.83) when compared to previously published iMGL using the same differentiation approach ^24^. In addition, we also verified that these cells did not express markers for other brain cell types (astrocytes, oligodendrocytes, and neurons; **Supplementary Figure 2 and Supplementary Table 2**).

### Brain Microglia vs iPSC-derived Microglia

We compared the transcriptomic profiles of our iMGL to both Fetal Brain and Adult Brain cell types ^17,25^. In both comparisons, we observed the highest correlation between iMGL and Fetal Brain Microglia (ρ= 0.711), and Brain Microglia (ρ= 0.637) compared to other brain cell types (**Table 2**). This data suggests these iMGL recapitulate well the transcriptomic profiles observed in brain microglia and are a good study model.

**Table 2:**
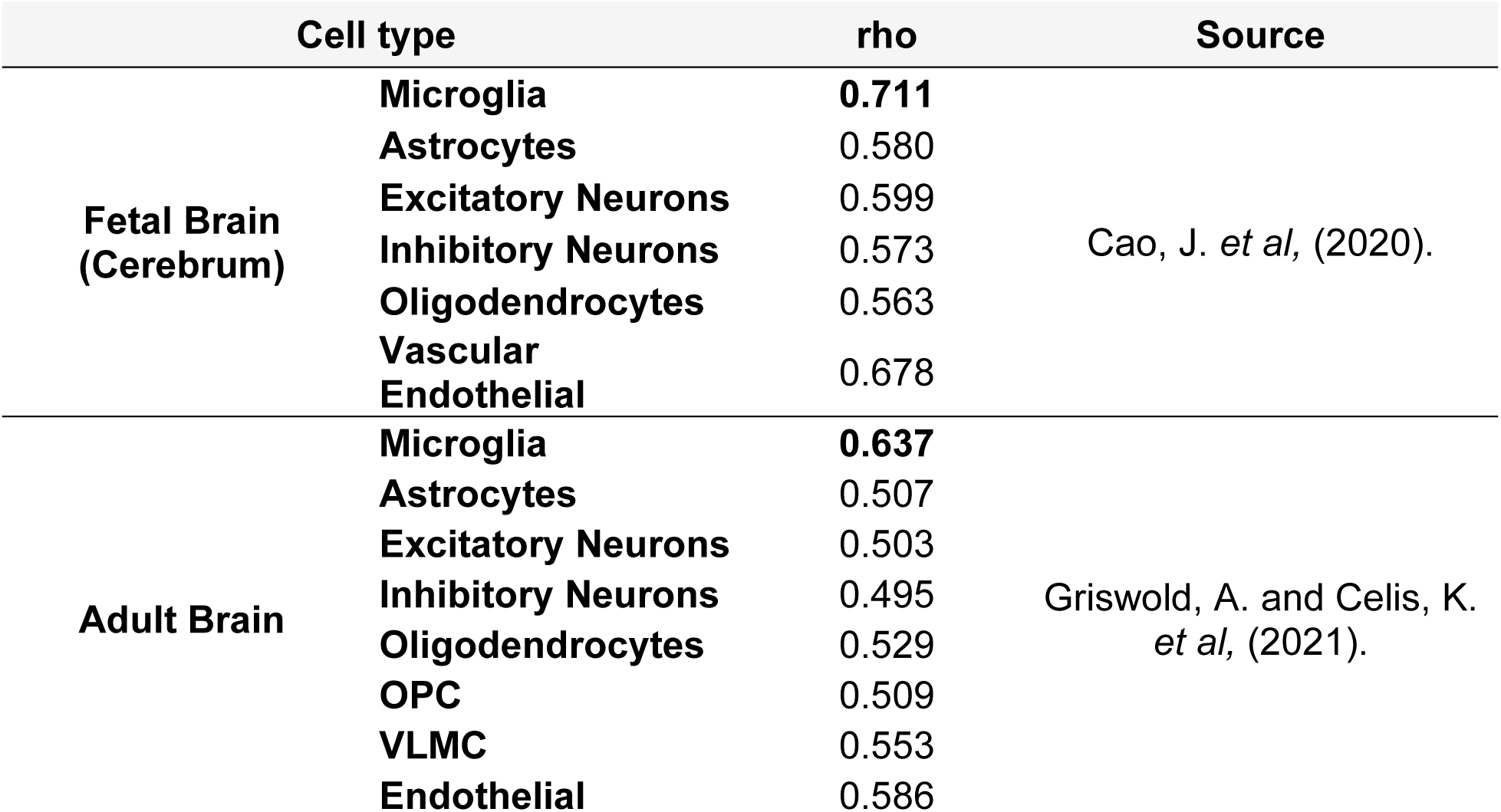
Correlation analysis between iMGL from our study and other cell types. Note that all thirteen iMGL lines were included for these comparisons and the p-value was below 2.2×10^-16^ for all comparisons. The Adult Brain data is derived from both African and European ancestry.

### Gene expression profiles across ancestries

We detected a total of 21,980 expressed genes across ancestries and performed differential expression pairwise comparisons between ancestries. In total, we observed 1,103 unique, differentially expressed genes (DEGs) between ancestries (FDR<0.05). Specifically, we identified 971 DEGs between Amerindian (AI) and AF, 320 between AI and EU ancestries, and 62 DEGs between African (AF) and Europeans (EU) (**Figure 1A and B; Supplementary Tables 3, 4, and 5**).

**Figure 1:**
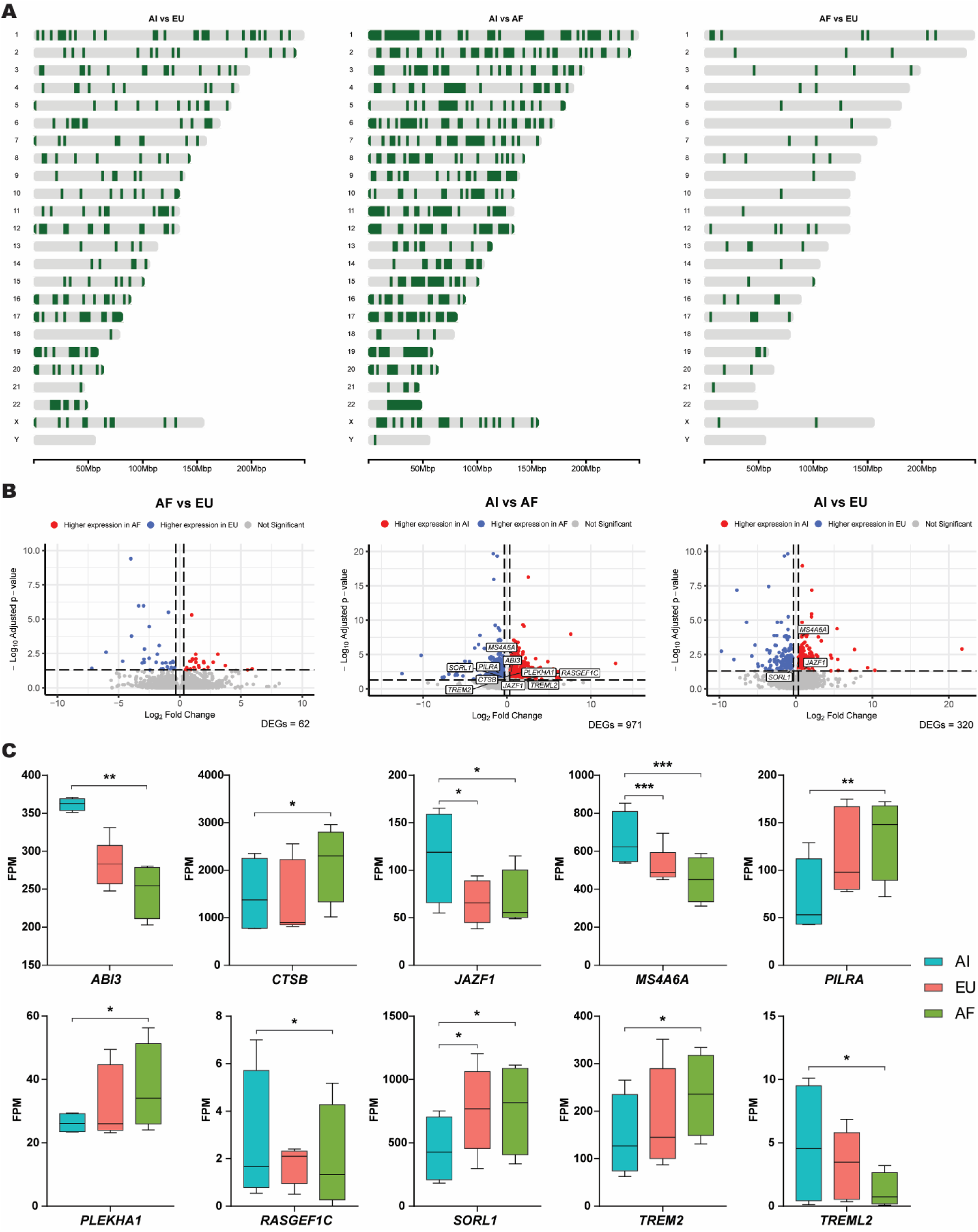
Gene expression across ancestries. **(A)** Chromosome Maps per pair-wise ancestral comparison demonstrating the distribution of differentially expressed genes (DEGs) genome-wide. The dark green color represents DEGs. **(B)** Volcano plots representing gene expression (Log_2_ Fold Change) per pair-wise comparison between ancestries (AF vs EU, AI vs AF, and AI vs EU). All 60,656 expressed variables are represented by the circles. The blue and red colored circles represent the genes that are differentially expressed (Fold Change cutoff of ± 1.25 and have an adjusted p-value (FDR)≤ 0.05). AD risk-modifying genes were highlighted in the white boxes. **(C)** Gene expression (FPM) of AD-related genes that were differentially expressed between ancestries. Box plots represent minimum to maximum FPM values and error bars denote the standard deviation. Asterisks denote adjusted p-value (FDR) with p≤ 0.05 (*), p≤ 0.01 (**), and p≤ 0.001 (***). FPM: Fragments per Million.

We focused on genes previously identified in AD GWAS studies ^5,26–30^. Of the 121 AD GWAS genes (**Supplementary Table 6**), we identified 10 DEGs between AI and AF (*ABI3, CTSB, JAZF1, MS4A6A, PILRA, PLEKHA1, RASGEF1C, SORL1, TREM2,* and *TREML2*) and 3 DEGs between AI and EU (*JAZF1, MS4A6A,* and *SORL1*). Despite our recent report on brain microglia of European and African ancestries ^17^, we did not observe differential expression for AD risk-modifying genes between AF and EU in our iPSC-derived microglia. We observed significantly higher gene expression in AI compared to AF for *ABI3, JAZF1* (also compared to EU), and *RASGEF1C,* while AF had significantly higher expression of *CTSB, PLEKHA1, SORL1,* and *TREM2* compared to AI. Lastly, we observed that EU express significantly higher amounts of *SORL1* compared to AI (**Figure 1C**).

### Chromatin accessibility across ancestries

We measured a total of 171,929 ATAC peaks for all ancestries and performed differential accessibility analysis genome wide. Overall, we observed 225 differentially accessible peaks (DAPs) linked to 208 unique, differentially accessible genes (DAGs) between AI and AF, 57 DAPs (55 DAGs) between AF and EU ancestries, and 53 DAPs (52 DAGs) between AI and EU (**Figure 2; Supplementary Tables 7, 8, and 9**). We observed an enrichment in DAPs between AI and EU in chromosome 17 (12.28%, Chi-square p-value=0.038) and chromosome 13 (7.02%, Chi-square p-value=0.041), which contain only ∼3% and ∼4% of the genome, respectively. Between AI and AF, we observed a significant enrichment in DAPs in chromosome 17 (9.78%, Chi-square p-value=0.004). Lastly, we observed that DAPs between AI and EU were enriched in chromosome 7 (5.66%, Chi-square p-value=0.031; **Figure 2A and Supplementary Table 10**). Overall, we observed that among all DAPs between all three ancestral comparisons, the DAPs lie primarily in intronic regions (∼28-39%) followed by distal intergenic (∼16-32%) and promoter regions (∼23-25%; **Figure 2B; Supplementary Table 11**). Interestingly, in the context of genes associated with AD, we only detected 2 DAGs (*PRDM7* and *SCIMP*) between AI and AF and 1 DAG between AI and EU (*PRDM7*) (**Figure 2C**).

**Figure 2:**
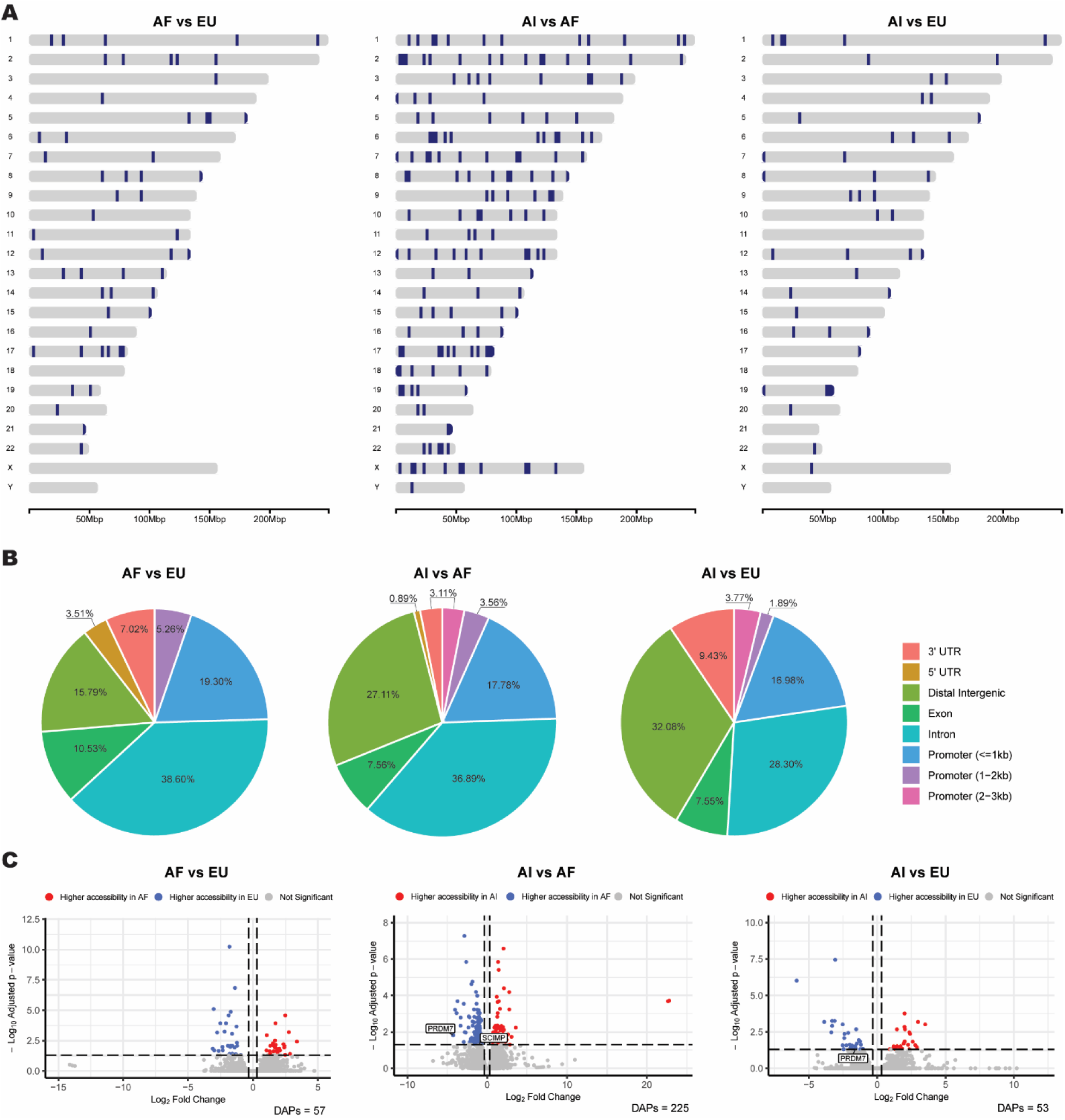
Chromatin accessibility across ancestries. **(A)** Chromosome Maps per pair-wise ancestral group comparison demonstrating the distribution of differentially accessible genes (DAGs) genome-wide. The dark blue color represents DAGs. **(B)** Pie Charts illustrate the regions of the genome in which the differentially accessible peaks lie for each of the ancestral comparisons. **(C)** Volcano plots representing chromatin accessible peaks (log_2_ Fold change) per pair-wise comparison between ancestries (AF vs EU, AI vs AF, and AI vs EU). All 171,929 peaks are represented by the circles. The blue and red colored circles represent the genes that are differentially accessible (Log_2_ Fold Change cutoff of ±0.322 and adjusted p-value (FDR)≤ 0.05. AD risk-modifying genes were highlighted in the white boxes.

We observed two DAPs in *PRDM7*: one in the proximal enhancer (Peak 1) and another in a distal enhancer (Peak 2; **Figure 3A**), according to ENCODE classification. Specifically, we observed that compared to AI, AF have significantly higher chromatin accessibility in peak 1 while EU have significantly higher accessibility in peak 2. Interestingly, contrary to other samples of the same ancestry group, we observed that sample 4 (AI) has chromatin accessibility in peak 1 while sample 6 (EU) presents visibly less accessibility in both peaks 1 and 2 (**Supplementary Figure 3**). We performed local ancestry (LA) analyses surrounding the *PRDM7* locus (± 500kb) to further investigate whether it could explain the differences in chromatin accessibility (**Supplementary Table 12**). We observed that samples 1-3 of AI global ancestry, have homozygote Amerindian LA for the *PRDM7* locus while sample 4 has African LA for both haplotypes in this locus aligning with the chromatin accessibility observations within the African global ancestry group. While this data suggests that the African LA of sample 4 in the *PRDM7* locus plays a role in and promotes chromatin accessibility, we did not observe any LA differences in the European global ancestry samples (all homozygote EU LA for this locus).

**Figure 3:**
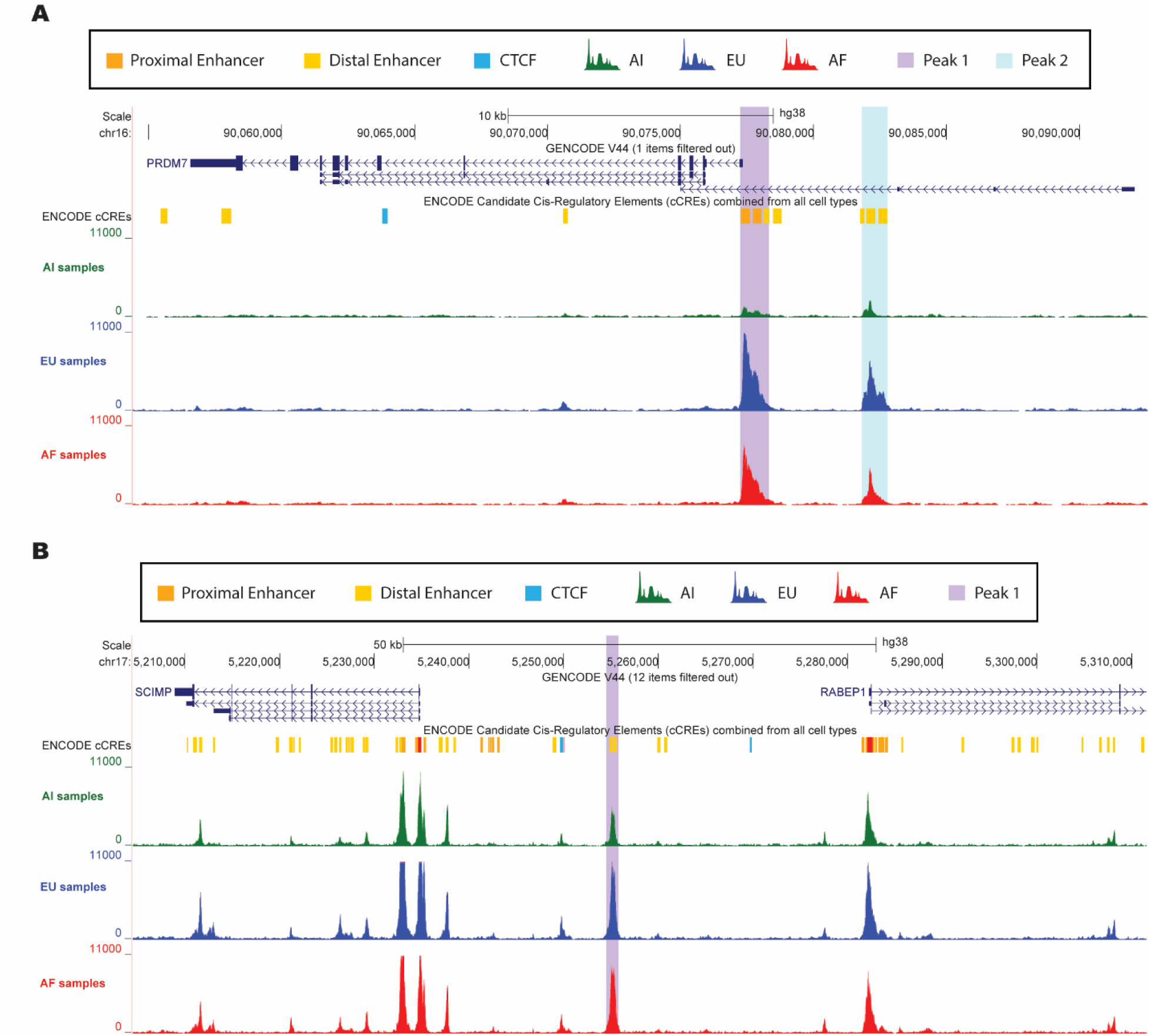
Differentially accessible peaks in AD-risk modifying genes across ancestries. **(A)** Differential chromatin accessible peaks in *PRDM7*. **(B)** Differential chromatin accessible peak in a distal intergenic enhancer of *SCIMP*. Note that the peaks represent merged data of all individuals within the same ancestry group.

In addition, we observed a DAP between AI and AF in a distal intergenic enhancer of *SCIMP* (∼20kb; **Figure 3B**). We did not observe LA differences within the same global ancestry group for the *SCIMP* locus (**Supplementary Table 12**) which could explain chromatin accessibility differences seen between global ancestry groups in this region (**Supplementary Figure 4**).

### Functional enrichment pathway analysis

To understand the functional mechanisms that might contribute to the differential AD risk across ancestries, we performed functional enrichment pathway analysis between the three ancestral groups using the g:Profiler tool in R. As expected, given the smaller number of DEGs between EU and AF, we only observed two significant functionally enriched pathways for these ancestries (**Supplementary Table 13**) and none have a known relation to AD. We observed that several DEGs across the other two ancestry group comparisons were involved in immune response, lysosomal activity, sterol and steroid biosynthesis and metabolism, cholesterol biosynthesis and metabolism, lipid transport and metabolism, and phagocytosis - all highly relevant processes in AD pathology (**Figure 4** and **Supplementary Tables 14 and 15**).

**Figure 4:**
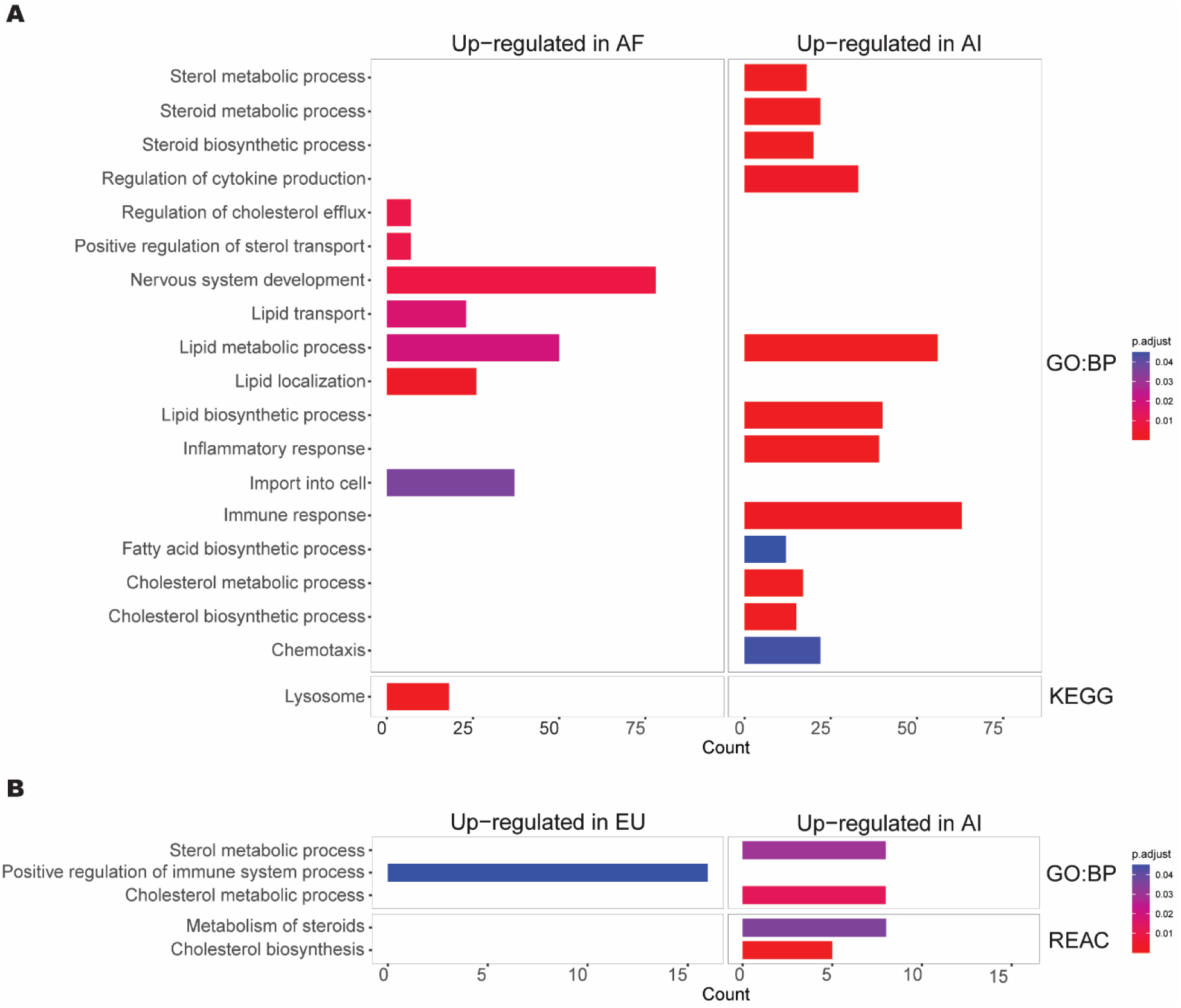
Functional enrichment pathway enrichment across ancestries relevant to AD. Pathway enrichment analyses between **(A)** AI and AF, and **(B)** AI and EU. See **Supplementary Tables 14 and 15**, respectively, for all significantly enriched pathways.

### Regulatory architecture in iPSC-derived Microglia

We studied the overlap between DAGs and DEGs to gain further insights into ancestry-specific regulatory mechanisms. Overall, we observed less than 2% shared DAGs and DEGs when comparing the ancestries (**Figure 5A** and **Supplementary Figure 5**). None of the overlapping DEGs and DAGs were from known AD GWAS genes. We observed that all overlapping DAGs and DEGs between AF and EU, and between AI and EU lay in promoter regions (**Supplementary Tables 16 and 17**, respectively) while there was a wider genomic distribution for those overlapping DAGs and DEGs between AI and AF (**Supplementary Table 18**).

**Figure 5:**
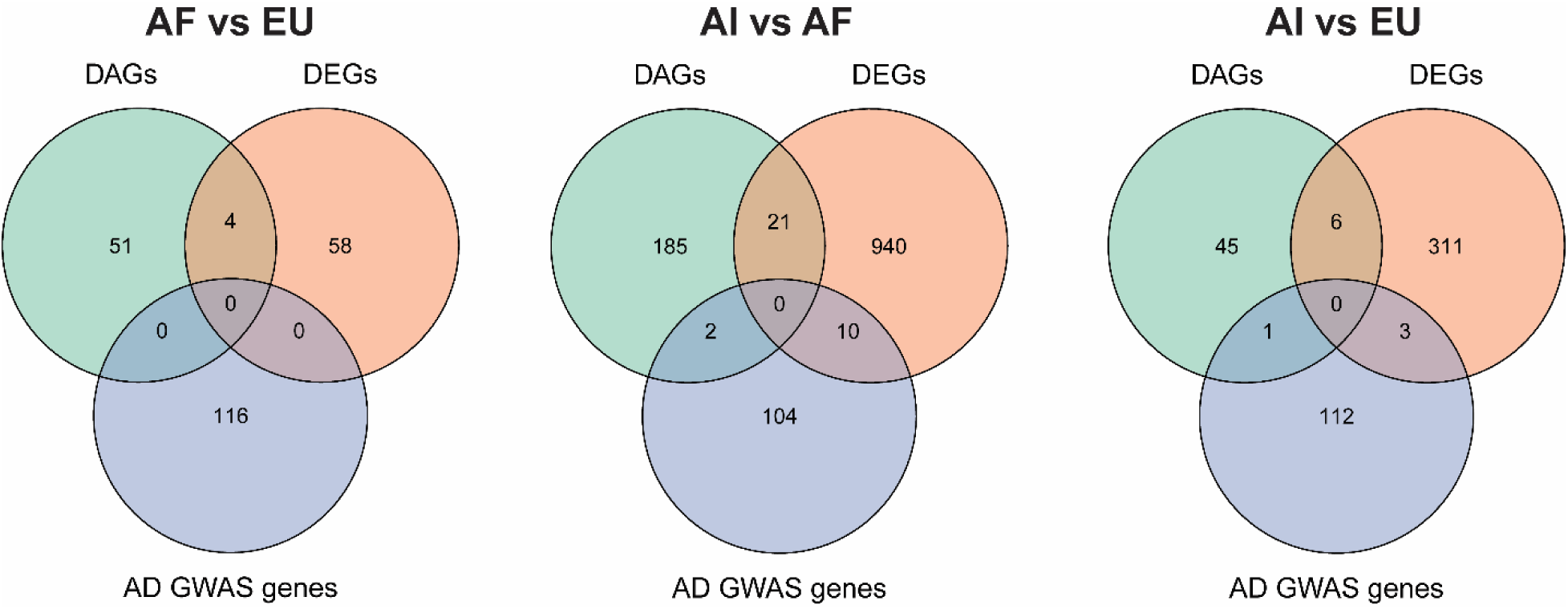
Overlap between differentially accessible ATAC-seq genes, differentially expressed RNA-seq genes, and AD GWAS genes between ancestry-group comparisons.

However, despite the small overlap between DAGs and DEGs with p-value≤ 0.05, we still observed a correlation between expression and chromatin accessibility in the promoter peaks (r= 0.53 (AF vs EU); r= 0.57 (AI vs EU); r= 0.47 (AI vs AF); **Supplementary Figure 6**).

### Regulatory differences specific to AD diagnosis, *APOE* genotype, and Sex

Between AD cases and controls, we performed differential expression analysis for 12 samples (the MCI sample was excluded from this analysis) and observed a total of 7 DEGs between non-cognitively impaired individuals and AD samples (**Supplementary Table 19**). None were previously identified as AD risk-modifying genes. Differential expression analysis between *APOEe3* and *APOEe4* homozygote carriers revealed 7 DEGs (**Supplementary Table 20**). Between the two analyses, we only found one DEG in common, high mobility group AT-hook 2 (*HMGA2*), which was overexpressed in AD and *APOEe4* carriers as compared to controls and *APOEe3* carriers (**Supplementary Figure 7**). The sex comparison revealed a total of 116 DEGs between Males and Females (**Supplementary Table 21**), none of which were AD risk-modifying genes or overlapped with any of the DEGs from the two aforementioned analyses. On the chromatin accessibility level, we only observed three DAPs/DAGs between *APOEe3* and *APOEe4* carriers (**Supplementary Table 22**), one DAP/DAG between cases and controls (**Supplementary Table 23**), and 136 DAPs between Males and Females (90 DAGs; **Supplementary Table 24**). None of these peaks have been previously connected to either AD or *APOE* genotype. Lastly, we observed an overlap between eleven sex-specific DEGs and DAGs, most of which are located in chromosomes X and Y.

### Ancestry-specific genetic regulatory architecture tool for other Neurological diseases

Despite the lack of ancestry-specific studies for other neurological diseases, ancestry might affect disease risk as observed in AD pathology. To demonstrate the importance of this GRA resource for the study of other neurological diseases in diverse ancestries, we compared both DEGs and DAGs identified for each of the ancestry comparison groups in our study with GWAS genes identified for Autism Spectrum Disorder (ASD) ^31–41^, Schizophrenia (SZ) ^42–57^, Bipolar disorder (BP) ^54,58–64^, Parkinson’s Disease (PD) ^65,66^, Multiple Sclerosis (MS) ^67,68^, Stroke ^69^, Coronary Artery Disease (CAD) ^70–76^, and Hyperlipidemia (HDL) ^77,78^ (**Figure 6**).

**Figure 6:**
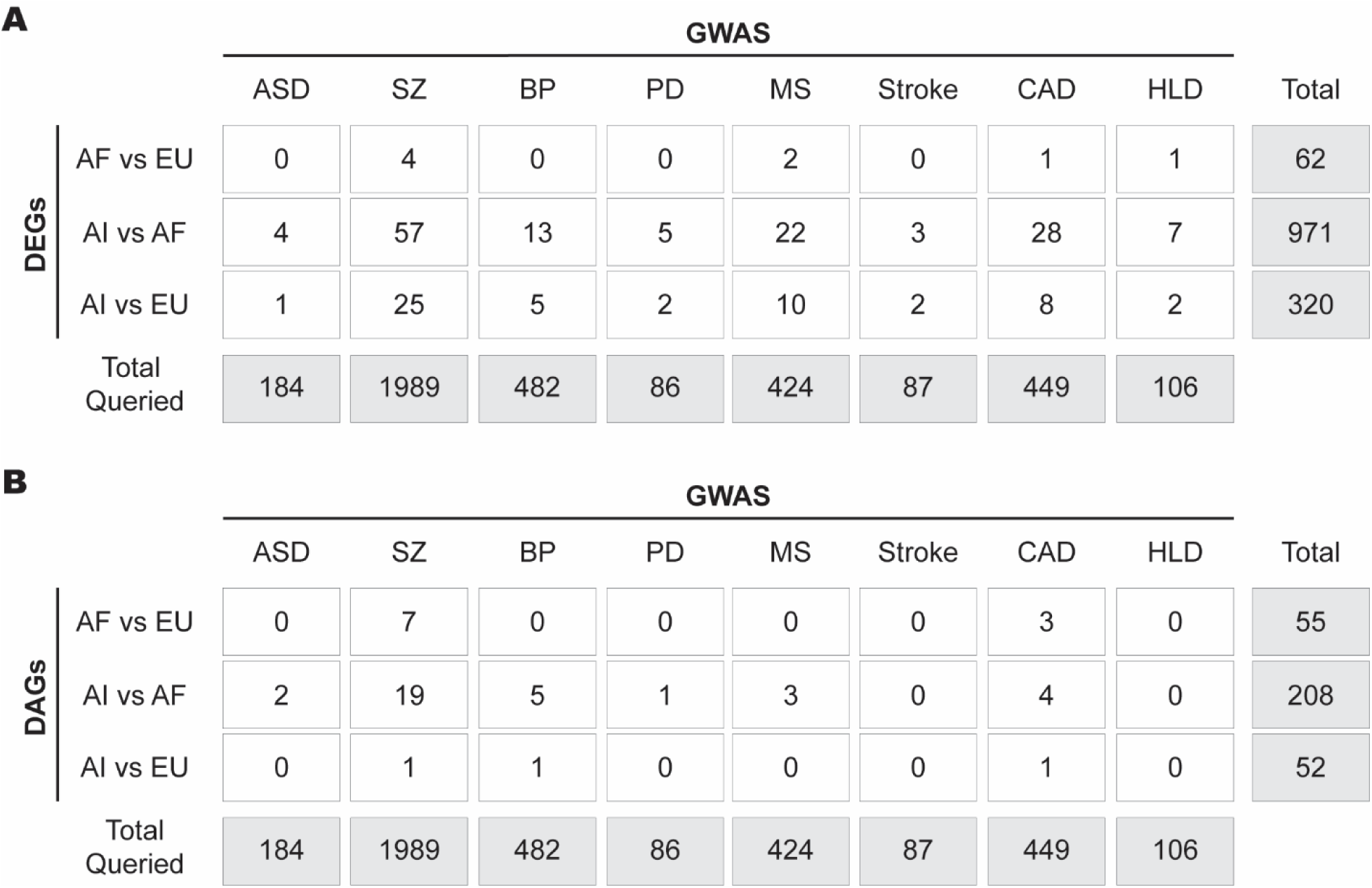
The genetic regulatory architecture in iMGL of diverse ancestries as a useful resource to study other neurological and associated diseases. We illustrate the overlap between ancestry-specific **(A)** DEGs and **(B)** DAGs from our study with previously identified GWAS genes for Autism Spectrum Disorder (ASD), Schizophrenia (SZ), Bipolar disorder (BP), Parkinson’s Disease (PD), Stroke, Multiple Sclerosis (MS), Coronary Artery Disease (CAD), and Hyperlipidemia (HLD). Gray boxes represent the total number of genes queried.

## Discussion

Recent studies have demonstrated that genetic disease associations differ in their strength and location between ancestries ^5,27,28,30^. As the majority of genetic associations are in non-coding regions, it is important to gain insight into the regulatory architecture of other ancestries besides European. Given the key role of microglia in AD pathology, we report, for the first time, epigenetic and disease-relevant differences between these ancestries in iMGL. While we have focused on AD, the microglial regulatory architecture presented here will be applicable to any study of the CNS.

Several known AD genes demonstrated ancestral expression differences in the microglia. One of these genes was ABI family member 3 (*ABI3)*, differentially expressed between AI and AF in this study and which has been previously found to be associated with AD in African American individuals ^79^. Studies have found that loss of *ABI3* function in mice was associated with Aβ-amyloidosis ^80^ and increased *ABI3* expression in microglia has been observed surrounding amyloid plaques in AD brain samples ^81^. Both studies hypothesize that *ABI3* expression plays a role in microglia migration in the central nervous system and affects disease progression in the absence of a functioning protein. We find that AF have on average the lowest expression of *ABI3*, compared to AI, supporting *ABI3* as an AD risk factor specifically in AF.

Another known AD gene, Cathepsin B (*CTSB)*, identified here as differentially expressed with higher expression levels in AF compared to AI, has been implicated as a major contributor to cognitive dysfunction and neuropathological changes, such as lysosomal dysfunction, cell death, and inflammatory responses ^82,83^. Interestingly, increased CTSB protein expression has been reported in AD patients compared to controls ^84–86^. It was also previously reported that *APOEe4* carriers of AF local ancestry expressed higher *CTSB* in brain microglia compared to those of EU local ancestry surrounding the *APOE* locus ^17^, similar to the trend observed in our dataset between AF and EU (**Figure 1C**). Again, this could suggest a larger role in AD risk for *CTSB* lying on AF local ancestry in African American individuals. Both of these differences were seen between AF and AI samples, which displayed the largest genomic differences between the three ancestries examined in this study. These are the two populations at either end of the migration spectrum for humans, implying these genetic ancestries had the longest time to evolve independently, creating ancestries who are the least related genetically.

In addition, even for genes without significant ancestral differences, the expression and accessibility data here can be useful for further understanding of the locus across population groups. For example, another AD-risk-modifying gene that showed differential gene expression is *MS4A6A.* This gene has been shown to be highly expressed in microglia ^87^ and it was previously reported that brain microglia of AF ancestry express less *MS4A6A* compared to those of EU ancestry ^17^. Despite not reaching significance, we did observe a similar trend towards less *MS4A6A* expression in AF iMGL compared to EU iMGL. *TREM2*, another well-known AD-GWAS gene, is primarily expressed in microglia and has been heavily implicated in AD progression ^88–91^. Interestingly, we found that AI cells express the lowest amount of *TREM2*. Data show that *TREM2* mRNA levels are associated with amyloid burden in cortical regions ^92^ and loss-of-function *TREM2* variants are associated with dementia ^93–95^, implying that the lower expression in AI microglia might impact AD risk in this ancestry due to reduced microglia functionality (Aβ-plaque clearance, *APOE*-mediated functions, immune modulation, and cell survival).

The iMGL lines used here varied not only in their genetic ancestry, but also in other variables such as sex, *APOE* genotype, and disease status which could complicate the interpretation of results. Therefore, we also performed differential expression analysis between Males and Females, AD vs controls, and *APOEe3* vs *APOEe4* carriers. Most of our AD patients were *APOEe4* homozygotes as at least 60% of AD patients carry the *APOEe4* allele. Despite observing a small number of DEGs between AD vs Controls and *APOE e3* vs *e4* carriers, we observed that *HMGA2,* a high-mobility protein that modulates transcription and chromatin condensation, was differentially expressed in both comparisons. Specifically, we observed higher gene expression in AD individuals and *APOEe4* carriers. Interestingly, silencing of *HMGA2* has been reported to lead to increased expression of the PI3K/AKT signaling pathway and improved memory and learning ability, reduced brain injury, and decreased oxidative stress and inflammatory reactions in mice ^96^. It was also recently reported that downregulation of *HMGA2* in AD patients was associated with increased lifespan ^97^. Thus, together with these findings, our results also suggest and support that increased *HMGA2* expression is a risk factor for AD.

We are often taught that chromatin accessibility is a key factor controlling gene expression. Comparing the significantly different changes in gene expression and chromatin accessibility between ancestries provides one opportunity to examine this relationship. Our differential analysis between ancestries revealed greater differences in gene expression (DEG) (approximately 0.3-4.4% of genes depending on the paired comparison) than in chromatin accessibility (DAP/DAG) (0.03-0.13%). This supports the growing understanding of the complexity of our cells in regulating gene expression and that transcription is a much more complex mechanism and higher accessibility is only one factor that could affect gene expression. For example, DNA sequence variability both at binding sites and distal eQTLs can complicate interpretation of the (dis)concordance between gene expression and chromatin accessibility changes. However, as expected, when expanding our sample size by using all our expression and accessibility data, we do find the expected moderate correlation between chromatin accessibility and expression (r=0.47 to 0.57).

iPSCs and derived cells have become important models for human brain disorders. We demonstrated that their transcriptome has a strong correlation with brain single nuclei RNAseq results ^17^. These iPSC-derived microglial cells were grown in the absence of other cell types and with a lack of environmental stressors. The complex gene regulatory networks operating in brain cells reflect the interplay of mostly invariable genetic factors with a dynamic exposome that includes chemical exposures, diet, and diverse stressors across the life course. One could postulate that microglia co-cultured with other CNS cell types or 3D organoids would feature cell-cell interactions that would provide an even stronger correlation with the brain transcriptome.

We did not observe any of the currently known African-specific AD GWAS genes ^5^ to be differentially expressed or accessible in the AF ancestry iMGL compared to the AI or EU ancestries. This could be explained by the fact that some of these genes were not expressed in iMGL and others had heterogenous expression levels between the limited number of individuals. The relatively small number of individuals included is the main limitation of this study. This is a general limitation of iPSC-derived cell studies which are expensive and time-consuming. Some of the differential findings reported here may reflect individual heterogeneity rather than ancestry generalizations. Additional iPSC-derived cell lines are needed to fully explore the regulatory architecture and to capture individual variability. Further genomic studies such as Hi-C will enhance these comparisons, particularly for specific genes of interest.

## Conclusions

Overall, we provide novel insights into the genetic regulatory architecture of microglia from three ancestry groups: Amerindian, African, and European. Transcriptional and architectural similarity was the most common finding, which is reassuring for future therapeutic interventions. We found a good correlation between the transcriptome of our iMGL and reported brain transcriptomes, as well as concordance for previously reported AD risk genes, supporting ancestral differences. These findings support the role of iMGL as a valuable model for human disease. Our data also supports a role for *HMGA2* expression in *APOEe4* carriers and AD risk. Lastly, this study provides a useful resource for the research community as it provides novel data on genome-wide regulatory architectures of diverse, understudied, genetic groups that could be applied to the study of other brain diseases, particularly those with high microglia involvement.

## Methods

### Sample collection

All samples of AI, EU, and AI cases and controls selected for this study were obtained from the John P. Hussman Institute for Human Genomics (HIHG) at the University of Miami Miller School of Medicine with the exception of the induced pluripotent stem cells derived from samples 7-9 which were obtained through ADRC from the University of California Irvine (UCI). All participants were ascertained using a protocol approved by the appropriate Institutional Review Board. This study received ethical approval from the University of Miami Institutional Review Board (approved protocol #20070307).

### Global ancestry ascertainment

We calculated the admixture proportions using a model-based clustering algorithm, as implemented in the ADMIXTURE software ^98^. A supervised ADMIXTURE analysis was performed at K = 4, incorporating four reference populations: 104 African, 84 European, 108 Amerindian, and 102 East Asian individuals from the Human Genome Diversity Project reference populations.

### Local ancestry ascertainment

To infer local ancestry, we first merged our dataset with the Human Genome Diversity Project reference panel, including European, African, and Amerindian reference populations ^99^. Next, we phased the combined data using SHAPEIT4 with default settings, referencing the 1000 Genomes Phase 3 reference panel ^100,101^. Finally, we estimated local ancestry at each genomic locus using RFMix v2 software ^102^.

### Whole Genome Sequencing (WGS)

DNA was extracted from all individual cell lines using the QIAamp DNA Blood Kit (QIAGEN, #51104) according to the manufacturer’s instructions. 1.5µg of DNA was submitted for WGS at the Center for Genome Technology (CGT) Sequencing Core at the HIHG using standard Illumina PCR-free library prep and sequencing protocols on the NovaSeq6000 followed by a bioinformatics pipeline incorporating the GATK Best Practices analysis recommendations ^103^. Individuals were screened for rare coding variants in seven AD-related genes nominated as likely causative by the ADSP Gene Verification Committee and variants in the promoter regions of the ten AD genes that had differential gene expression (**Supplementary Table 1**).

### Induced pluripotent stem cell generation

Peripheral blood mononuclear cells (PBMCs) were isolated from whole blood using SepMate-50 tubes with Lymphoprep (STEMCELL Technologies, #85450 and #07801) through density-gradient centrifugation according to the manufacturer’s instructions. PBMCs were reprogrammed into induced pluripotent stem cells (iPSCs) using CTS™ CytoTune™-iPS 2.1 Sendai Reprogramming Kit (Invitrogen, #A34546) according to the manufacturer’s instructions. Reprogrammed cells were tested for Sendai Virus absence, trilineage differentiation capability, immunocytochemistry, STR profiling, karyotyping, and mycoplasma testing as previously described ^104^. PBMC isolation and reprogramming was performed at the Hussman Institute for Human Genomics (HIHG) Induced Pluripotent Stem Cell (iPSC) Core at the University of Miami. Validation analyses were performed by the HIHG-iPSC Core and WiCell.

### Differentiation of iPSCs to Microglia

iPSCs were differentiated into hematopoietic progenitor cells (HPCs) and subsequently into Microglia (MGL) as previously described ^24^ with minor modifications.

In brief, feeder-free iPSCs were cultured and expanded in StemFlex medium (Gibco^TM^, #A3349401) in vitronectin (10µg/ml, Gibco^TM^, #A31804) coated cell culture-treated plates. On day −1, iPSCs were passaged with 0.5M EDTA onto Matrigel-coated (Corning, #354277) 12-well plates at a density of 10-20 aggregates/cm^2^ (>50µm in size). On day 0, if 4-10 colonies/cm^2^ adhered, the StemFlex medium was replaced with 1ml/well of HPC medium A (Basal medium with supplement A (1:200), STEMCELL Technologies, #05310). Half-medium change was carried out 48 hours later. On day 3, HPC medium A was replaced in full by medium B (Basal medium with supplement B at 1:200). Half-medium changes of medium B were performed on days 5, 7, and 10. HPCs were harvested on day 12.

On day 0 of microglia differentiation (day 12 of HPC differentiation), HPCs were plated at 22,000 cells/cm^2^ onto a Matrigel-coated 6-well plate containing 2ml of Microglia differentiation medium (Basal Medium with supplement 1 and 2 at 1:9 and 1:225, respectively; STEMCELL Technologies, #100-0019). Cells were supplemented with fresh half-medium every other day from day 0 to day 10. On day 12, cells were collected and centrifuged at 300 x g for 5 minutes. The cell pellet was resuspended in 2ml/well of fresh Microglia differentiation medium and transferred to a freshly Matrigel-coated 6-well plate. Cells were supplemented with 1ml of media every second day until day 22. Microglia cells were collected, resuspended in 2ml of Microglia maturation medium (Basal Medium with supplement 1 (1:9), and 2 and 3 (1:225); STEMCELL Technologies, #100-0020), and re-plated for assays into new Matrigel-coated 6-well plates. Lastly, on day 26, microglia were harvested for immunocytochemistry (ICC), bulk RNA-, and ATAC-sequencing.

### RNA isolation and sequencing

Total RNA was isolated from 1 million microglial cells per cell line using the RNeasy Mini kit (QIAGEN, #74104) according to the manufacturer’s instructions. Suspension cells were collected and centrifuged for 5 minutes at 300 x g. 600µl of RLT buffer (including β-Mercaptoethanol at 1/100) was used to collect semi-attached microglia and subsequently resuspend the cell pellet from the previous step. Cells were briefly vortexed for 1 minute and homogenized by loading the lysate into a QIAshredder spin column (QIAGEN, #79656) and centrifuging for 2 minutes at full speed. The homogenized lysate was resuspended in 1 volume of 70% ethanol and transferred to a RNeasy spin column and centrifuged for 30 seconds at 8,000 x g. 350µl of Buffer RW1 was added to the same spin column and centrifuged for 15 seconds at 8,000 x g. Following this, 80µl of DNAse I incubation mix (70µl of RDD buffer and 10µl of DNAse I, QIAGEN, #79254) were added to the spin column and incubated at RT for 15 minutes. Buffer RW2 (350µl) was transferred to the spin column and centrifuged for 15 seconds at 8,000 x g. 500µl of RPE buffer were loaded into the column followed by a centrifugation step of 30 seconds at 8,000 x g. The previous step was repeated once again but centrifuged for 2 minutes at 8,000 x g to ensure all residual ethanol was removed. The RNeasy spin column was transferred to a new 1.5ml collection tube and 30µl of RNAse-free water were added to the column to elute the bound RNA. Lastly, the spin column was centrifuged at 8,000 x g for 1 minute and then stored at −80°C until further used. The RNA concentration and quality were assessed using the Agilent Tapestation (Agilent Technologies) to determine the RNA integrity number (RIN).

### Bulk RNA sequencing

RNA libraries were prepared at the John P. Hussman Institute for Human Genomics Center for Genome Technology (University of Miami, FL) from ribodepleted total RNA. In brief, total RNA was prepared with the TECAN Universal Plus Total RNA-seq with NuQuant^®^ Human AnyDeplete according to the manufacturer’s instructions, using 60ng via QuBit and 16 PCR cycles. The normalized libraries were sequenced as paired end 100bp reactions targeting 30 million reads/sample on the Illumina NovaSeq 6000 (Illumina, CA). The raw FASTQ files were processed through an in-house bioinformatics pipeline including adapter trimming by TrimGalore (v0.6.10) (https://github.com/FelixKrueger/TrimGalore), alignment to the GRCh38 human reference genome with STAR (v2.5.0a) ^105^, and gene counts quantified against the GENCODEv35 gene annotation release using the GeneCounts module implemented in STAR.

### Bulk ATAC-sequencing

Cultured cells were treated with DNase I (200U/mL; QIAGEN, #79254) at 37°C for 30 minutes. The treated cells were then harvested and pelleted at 400 x g for 5 minutes at 4°C. The cell pellet was carefully washed in cold 1x PBS. The cells were re-pelleted as described before and then lysed in 100µl of lysis buffer (10mM Tris-HCl pH 7.4, 10mM NaCl, 3mM MgCl_2_, 0.1% NP-40, 0.1% Tween-20, and 0.01% Digitonin) on ice for 5 minutes. Next, the lysed microglia were washed in 1ml of wash buffer (10mM Tris-HCl pH 7.4, 10mM NaCl, 3mM MgCl_2_, and 0.1% Tween-20) and 100,000 nuclei were pelleted at 500 x g for 10 minutes at 4°C. The nuclei were incubated at 37°C for 30 minutes at 1,000rpm in 100µl of Transposition mix (2x Tagment DNA Buffer, 1x PBS, 0.1% v/v Tween-20, 0.01% v/v Digitonin, and 5µl of Tagment DNA Enzyme 1). The transposed DNA was purified using the MinElute PCR Purification kit (QIAGEN, #28004) and eluted in 10µl of Elution Buffer. The purified transposed DNA was combined with 25µM of Custom Adapter 1 (no primer mix), 25µM of Custom Adapter 2 (barcode), and NEBNext High-Fidelity 2x PCR Master Mix and ran on a thermocycler with the following conditions: 72°C for 3 minutes, 98°C for 30 seconds, and 5 cycles of 98°C for 30 seconds, 63°C for 30 seconds, and 72°C for 1 minute. The additional number of cycles required was determined as described in ^106^ and ran with the same conditions abovementioned. The amplified libraries were purified with the MinElute PCR Purification kit and eluted in 20µl of Nuclease-free water. Library traces were assessed by the Agilent Tapestation and when necessary, size selection purification was carried out using the AMPure XP beads (Beckman Coulter, #A63880) according to the manufacturer’s instructions. See **Supplementary Table 26** for full adapter sequences. Libraries were sequenced in paired end 100bp reactions targeting 30 million reads/sample on the Illumina NovaSeq 6000. The ATAC-seq data were preprocessed (trimmed, aligned, filtered, and quality-controlled) and analyzed using an adapted version of the ENCODE ATAC-seq pipeline. In brief, adapters and poor-quality bases were trimmed using TrimGalore (v0.6.10) (https://github.com/FelixKrueger/TrimGalore). Reads were aligned to the CRCh38 human reference genome with bowtie (v2.2.2) ^107^, duplicates marked with Picard (v2.1.1) (https://broadinstitute.github.io/picard/), and peaks called using MACS2 (v2.2.7.1) ^107^. Peaks were merged across all samples using an overlapping peak/union strategy to obtain a list of peaks across all samples. Counts per peak were calculated from individual aligned BAM files using htseq-count (v1.99.2) using the un-stranded option.

### Differential expression and accessibility analyses

Differential expression and accessibility analyses were carried out across the different ancestral populations using DESeq2 (version 3.17) package ^108^ in R language environment (version 4.2.1). We used DESeq2 default parameters and controlled for batch differences (design = ∼batch + ancestry). Three contrasts were run: AF vs EU, AI vs AF, and AI vs EU. Genes that were significantly expressed and/or accessible were identified with an FDR adjusted p-value of <0.05.

### Functional enrichment pathway analysis

Functional enrichment analysis was done with the R library gprofiler2 ^109^. We extracted gene symbols of DEG between ancestries (FDR adjusted p-value of <0.05), and the function *gost* was used to perform the gene set enrichment analysis for each ancestry comparison using the Gene Ontology, KEGG pathways, and REACTOME databases. Multiple comparison correction of enrichment scores was done with the ‘gSCS’ method. Pathways were considered significant if p-adj < 0.05. Results were manually curated to show known pathways related to AD pathogenesis, and the corresponding full lists of enriched terms are described in **Supplementary Tables 13, 14, and 15**.

### ATAC peak annotation

The function *annotatePeak* from Chipseeker R library ^110^ was used to annotate peaks with the nearest gene and genomic region. The annotation was done at the transcript level using the GENCODE V44 database. The distance of ±3 kb from the transcription start sites (TSS) was used to assign a peak to a gene promoter-TSS, and the following priority was defined for annotation: "Promoter", "5UTR", "3UTR", "Exon", "Intron", "Downstream", "Intergenic”.

### Immunocytochemistry (ICC) and fluorescence imaging

Cultured microglia cells were fixed with 4% formaldehyde for 15 minutes at RT and washed with 1x PBS. Cells were permeabilized for 10 mins with PBS-T solution (0.1% Triton X and 1x PBS). The microglia cells were then incubated in blocking buffer (1x PBS and 5% normal donkey serum) for 1 hour at RT. The blocking buffer was removed and incubated in the primary antibody solution (1% donkey serum, 0.1% Tween-20, 0.01% Sodium Azide, and target primary antibody) at 4°C overnight. The following day, the primary antibody solution was removed, and the cells were washed three times with 1x PBS. Following this, the secondary antibody solution (1% donkey serum, 0.1% Tween-20, 0.01% Sodium Azide, and secondary antibody) were added to each well, and cells were incubated for 1 hour at RT in the dark. Lastly, the secondary antibody solution was removed, and cells were washed thrice with PBS. The cells were washed with 1x PBS and incubated with DAPI (NucBlue Fixed Cell Stain). Images were acquired using a Keyence Microscope BZ-X800. See Supplementary Table 25 for details on all antibodies used for ICC analysis.

### Correlation analyses between differential expression and differential accessibility

Pearson correlation (r) was used to evaluate the relationship between gene expression and corresponding promoter accessibility. First, DEGs between ancestries with |log_2_(FoldChange)|≥ 1 and adjusted p-values ≤ 0.1 were considered for the analysis. Then, promoter peaks (distance of ±3 kb from TSS) annotated to those DEGs were considered for correlation analysis.

### Correlation analyses between iPSC-derived Microglia and other cell types

Correlation analyses between iMGL and Brain cell types were performed using Spearman correlation analyses. Specifically, we calculated the average expression of all thirteen iPSC-derived Microglia (iMGL) cell lines included in this study for each gene. Note that genes with an expression value of 0 were excluded as well as sex-related (Chromosomes X and Y) and mitochondrial genes. Following this, genes were ranked in descending order by expression level for both iMGL and brain cell types, and only genes present in both comparison datasets were included in the Spearman correlation test.

## Data Availability

All data generated or analyzed during this study are included in this published article and its supplementary information files. Sequencing files can be requested to the corresponding author.

## Acknowledgments

This study was supported by the National Institute on Aging (grant numbers U01-AGO72579, RF1-AGO59018, U01-AG066767, U01-AG052410, R56-AG072547, R01-AG070864).

We acknowledge the Center for Genome Technology (CGT) from the John P. Hussman Institute for Human Genomics (HIHG) from the University of Miami, Miller School of Medicine for the genomic and data analyses. We thank Dr. Lily Wang for meaningful data discussions and express our gratitude to the numerous participants, researchers, and staff involved for their invaluable contributions to the present study.

## Author contributions

S.M., D.M.D, A.J.G., J.I.Y., and J.M.V. conceptualized the project and planned experiments. S.M. performed the experiments. S.M., L.B.N., and A.J.G. analyzed the data. A.M.R., and L.C., contributed to the performance of experiments. B.DR. assisted with iPSC reprogramming. J.R. and F.R. performed local and global ancestry analyses. D.V.B. and L.B.N. performed bioinformatic analyses. K.H.-N., P.W., L.A., T.S., P.M., M.I.-M., S.T., G.B., M.C.-O., B.F.-A., and M.A.P.-V. contributed to sample collection and processing. S.M., L.B.N., A.M.R., B.DR., K.N., L.W., D.M.D., F.R., A.J.G., J.I.Y., and J.M.V. discussed the data results. S.M., L.B.N., F.R., A.J.G., D.M.D., J.I.Y, J.M.V. wrote the manuscript. All authors read and approved the final manuscript.

