## Supplementary Figures for "Gene expression and chromatin accessibility comparison in iPSC-derived microglia in African, European, and Amerindian genomes in Alzheimer’s patients and controls"

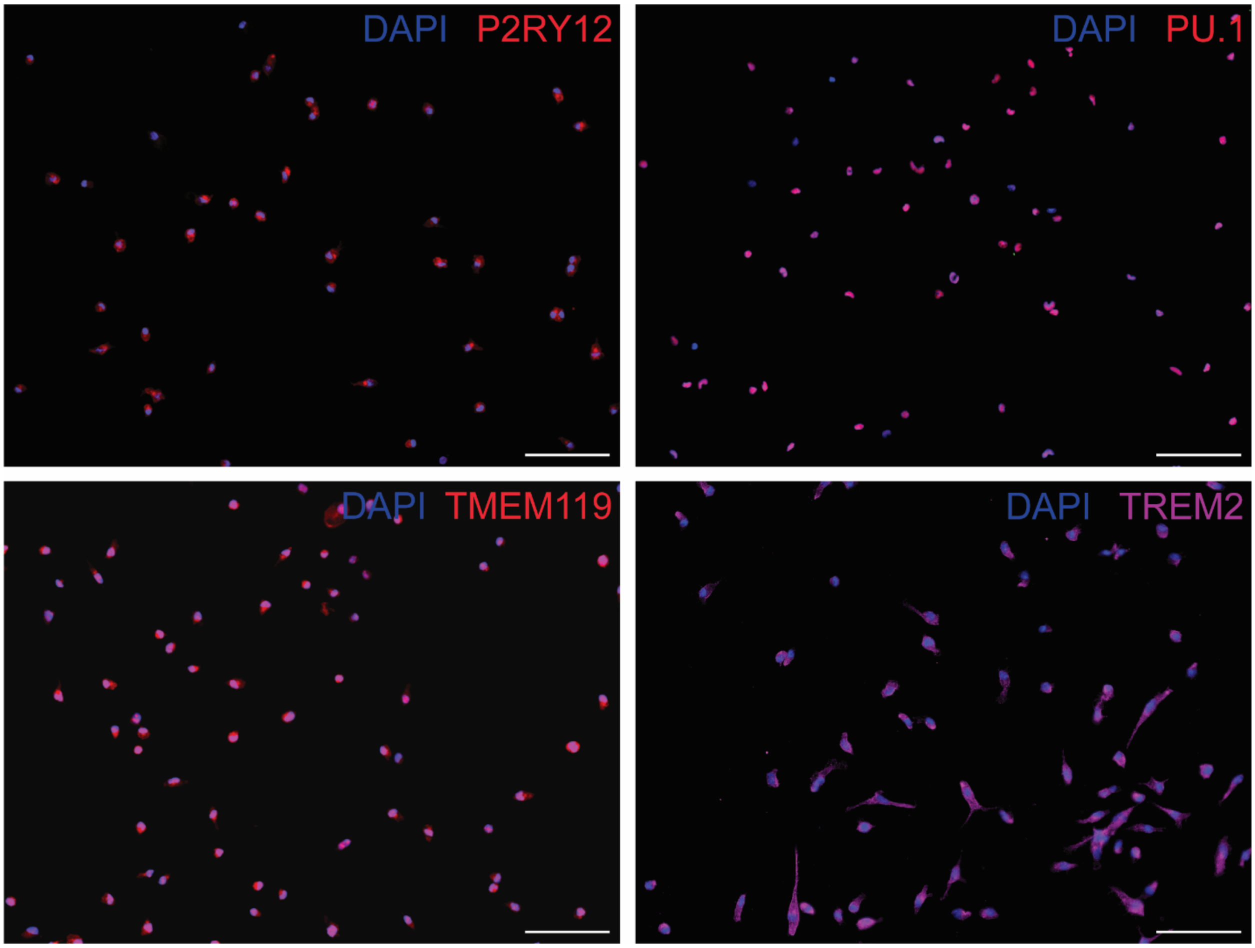


**Supplementary Figure 1:** iPSC-derived Microglia validation. Immunocytochemistry validation for microglia lineage specific markers: P2RY12, PU.1, TMEM119, and TREM2. These representative pictures illustrate overlay of single-staining images with the blue staining representing DAPI (cell nuclei). Scale bar: 100µm.

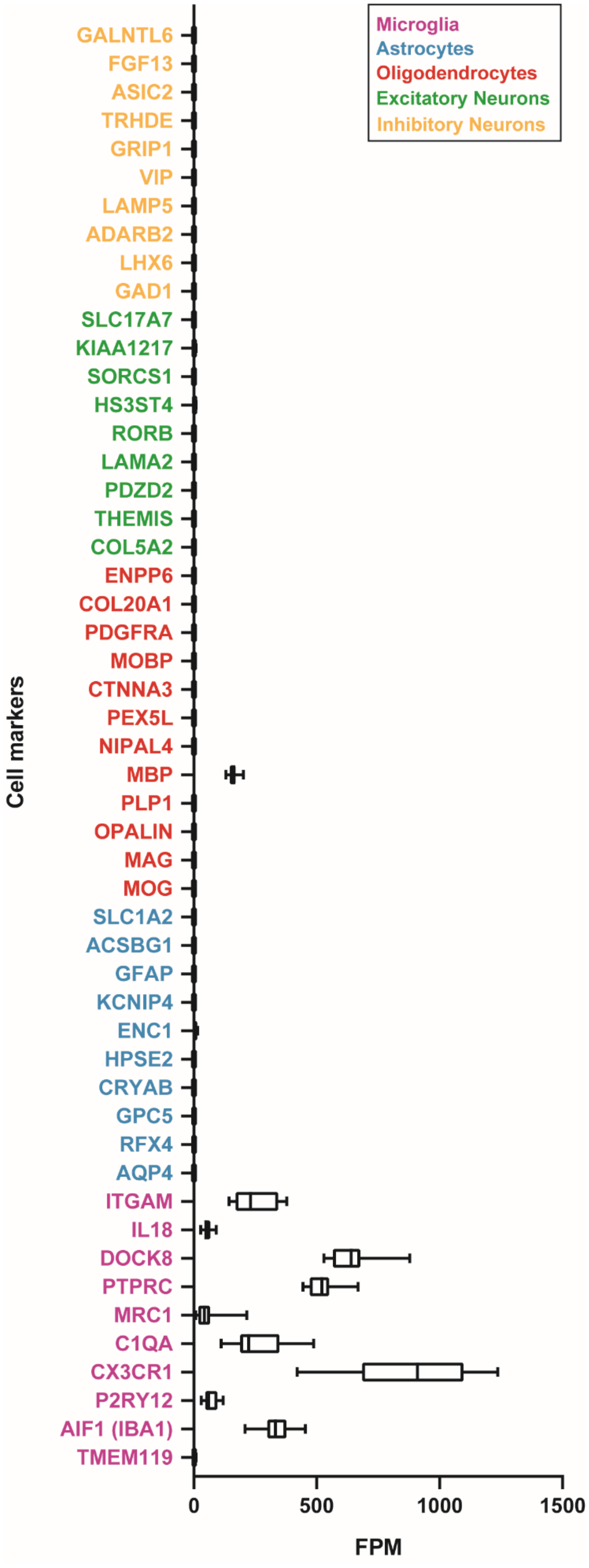


**Supplementary Figure 2:** Expression data for cell type-specific markers. The boxplots represent the expression data (FPM) of all 13 iPSC-derived Microglia cell lines. Error bars represent 95% Confidence Interval.

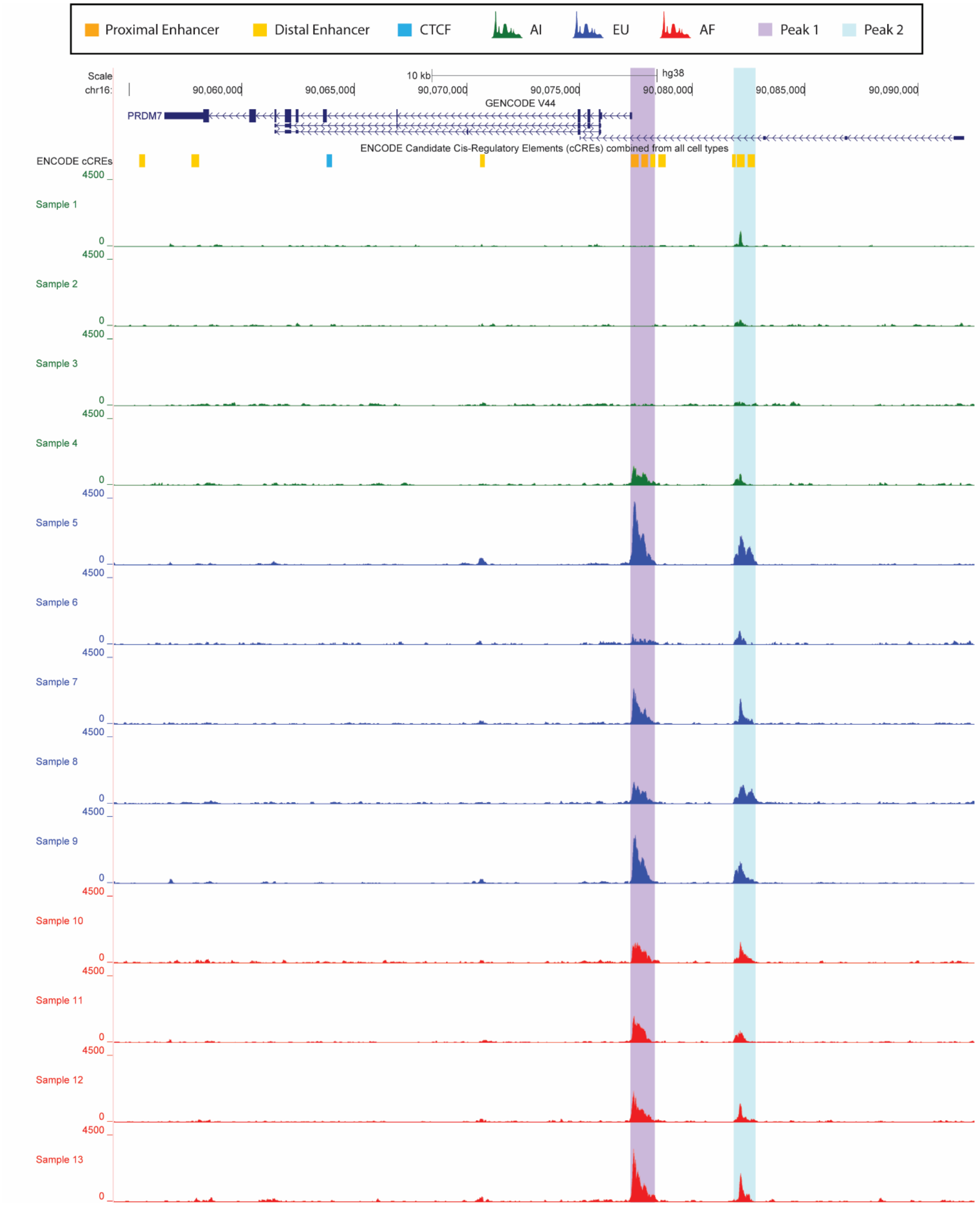


**Supplementary Figure 3:** Differentially accessible peaks in *PRDM7*. Peak 1 (purple highlight) overlaps a proximal enhancer of *PRDM7* and is differentially accessible between Amerindian (AI) and African (AF) ancestries. Peak 2 (blue highlight) overlaps a distal enhancer of *PRDM7* and is differentially accessible between AI and European (EU) ancestries.

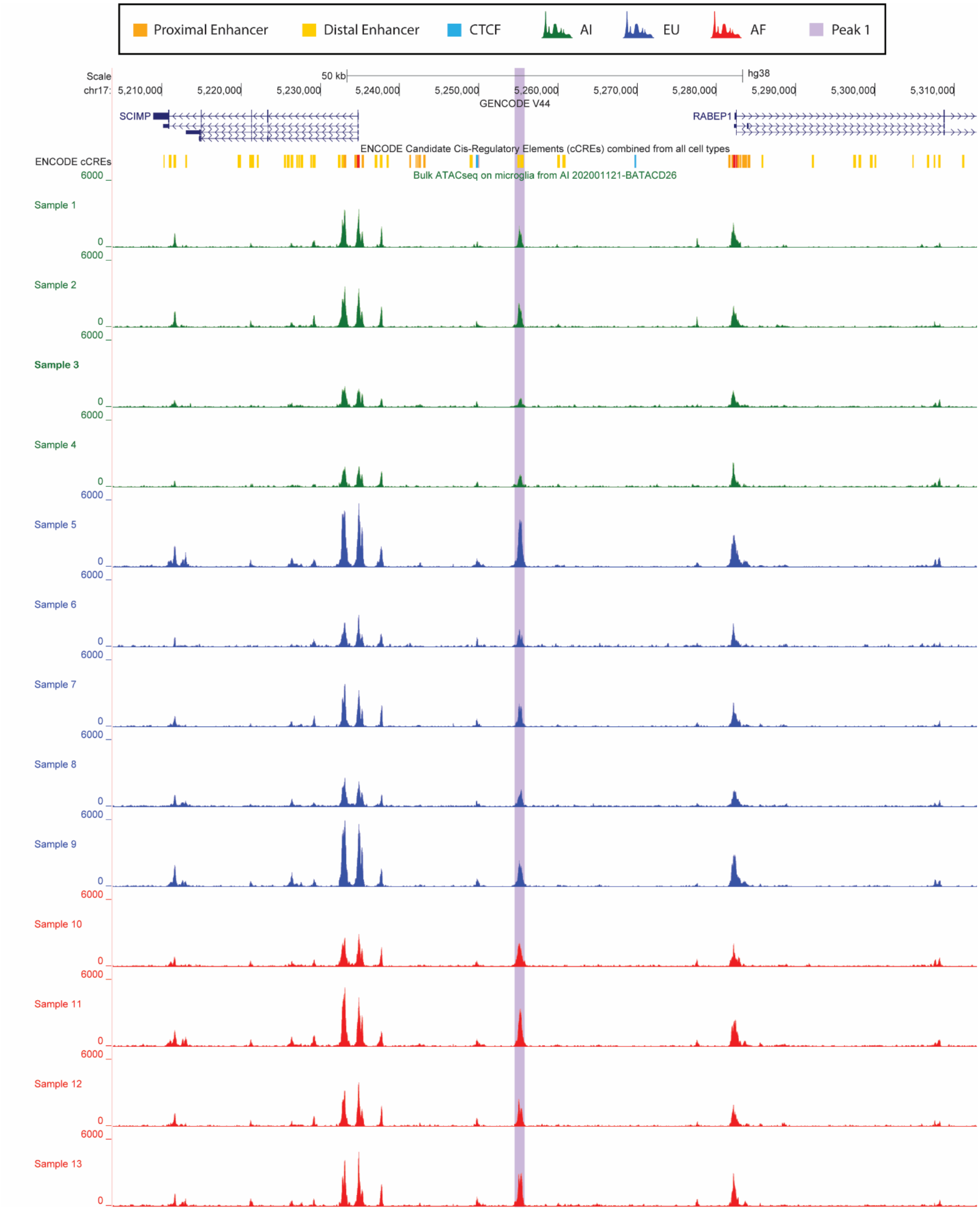


**Supplementary Figure 4:** Differentially accessible peak in a distal intergenic enhancer of *SCIMP.* The peak area highlighted in purple is differentially accessible between AI and AF and overlaps a distal enhancer of *SCIMP.* This peak lies in an intergenic region between *SCIMP* and *RABEP1* but is closer to the transcription start site (TSS) of *SCIMP.*

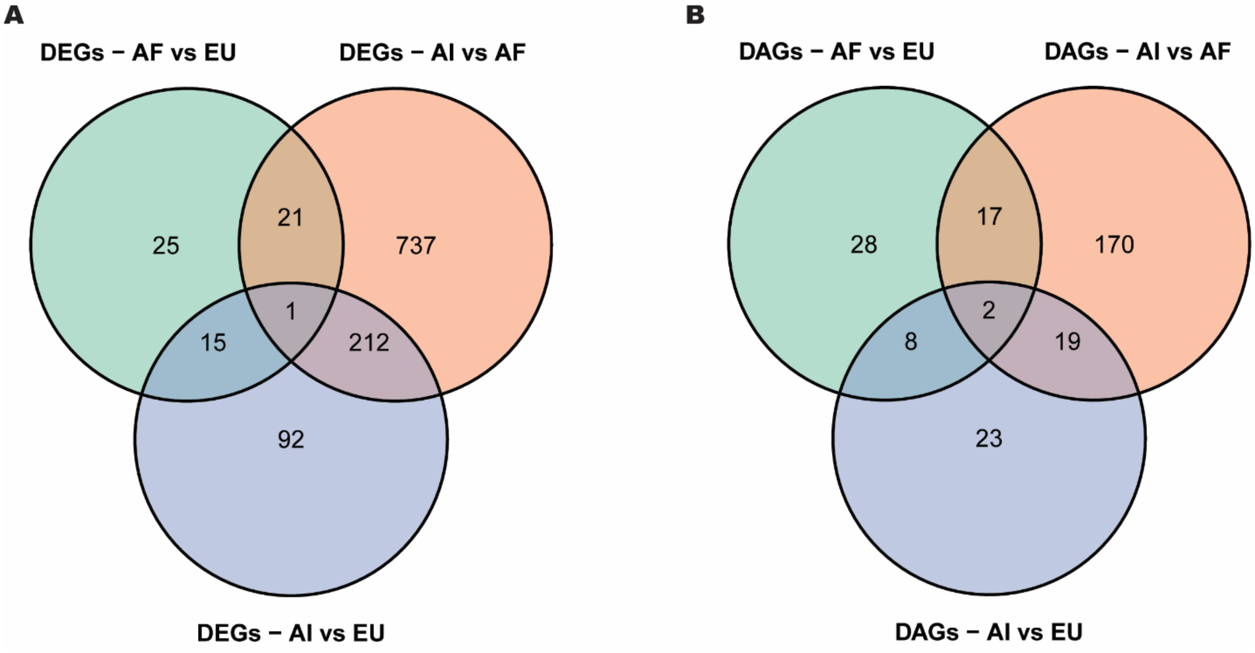


**Supplementary Figure 5:** Genetic regulatory architecture differences across ancestries. **(A)** Differential expressed genes (DEGs) and **(B)** differential accessible genes (DAGs) in the three ancestry group comparisons.

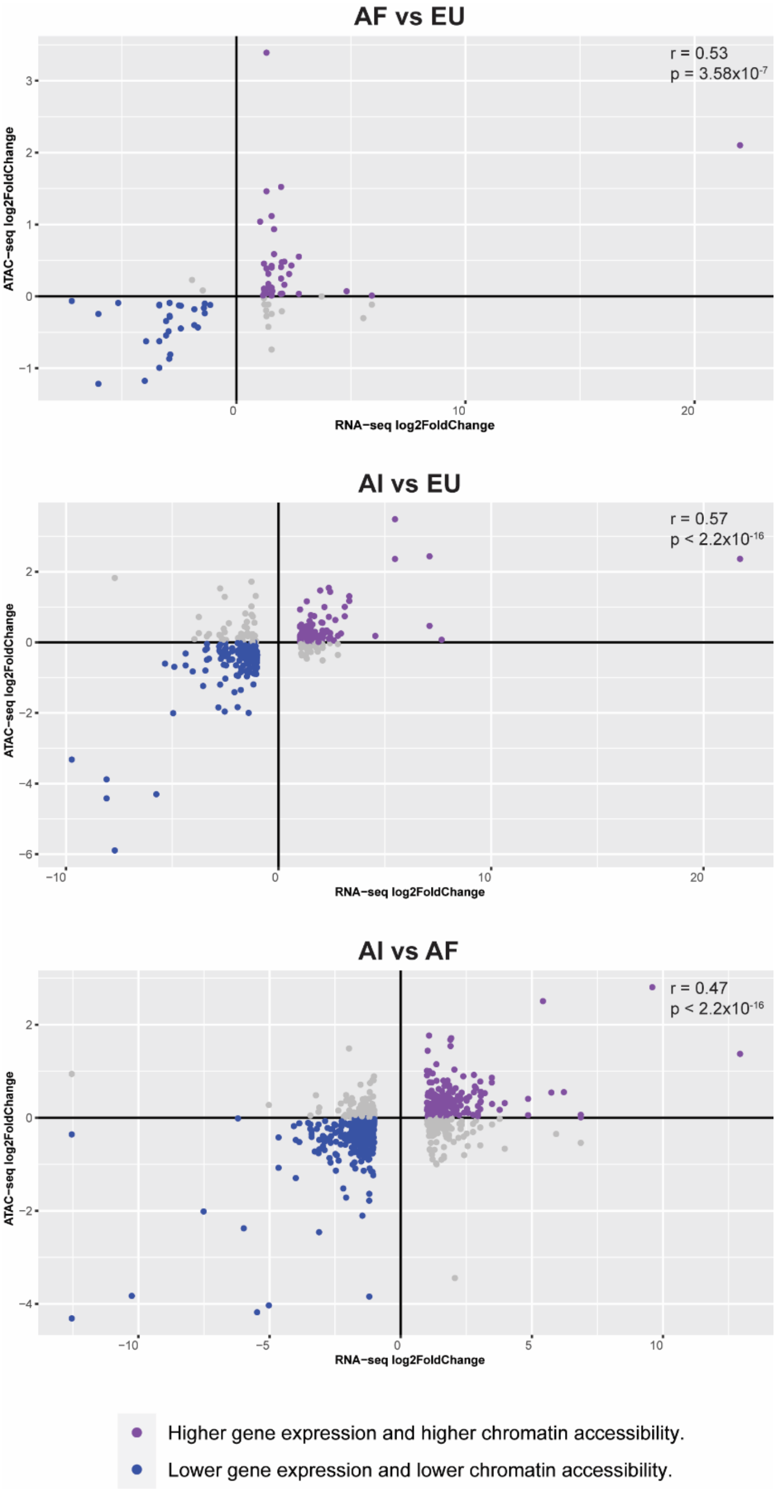


**Supplementary Figure 6:** Correlation between chromatin accessibility and gene expression between ancestry comparisons. Purple circles represent differentially expressed genes with positive log2 Fold Change and also positive log2 Fold Change for chromatin (not necessarily significant differentially accessible chromatin peaks). Blue circles represent differentially expressed genes with negative log2 Fold Change and also negative log2 Fold Change for chromatin (not necessarily significant differentially accessible chromatin peaks).

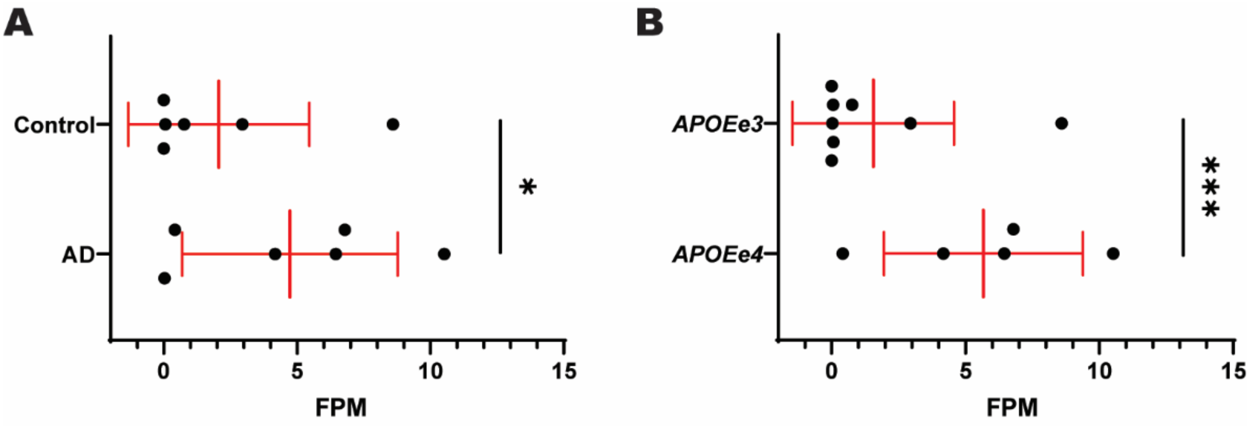


**Supplementary Figure 7:** Differentially gene expression of *HMGA2*. Gene expression between **(A)** Control and AD individuals and **(B)** *APOEe3* and *APOEe4* carriers. Note that control individuals (n=6) are *APOEe3* carriers (n=8) while AD individuals (n=6) are mostly *APOEe4* carriers (n=5). Mean and standard deviation are represented in red. Individual data points for expression of *HMGA2* are represented by circles. FPMs: Fragments per million. Asterisks denote adjusted p-value (FDR) with p≤ 0.05 (*), and p≤ 0.001 (***).
